# The structural determinants of intra-protein compensatory substitutions

**DOI:** 10.1101/2021.11.11.468231

**Authors:** Shilpi Chaurasia, Julien Y. Dutheil

## Abstract

Compensating substitutions happen when one mutation is advantageously selected because it restores the loss of fitness induced by a previous deleterious mutation. How frequent such mutations occur in evolution and what is the structural and functional context permitting their emergence remain open questions. We built an atlas of intra-protein compensatory substitutions using a phylogenetic approach and a dataset of 1,630 bacterial protein families for which high-quality sequence alignments and experimentally derived protein structures were available. We identified more than 51,000 positions coevolving by the mean of predicted compensatory mutations. Using the evolutionary and structural properties of the analyzed positions, we demonstrate that compensatory mutations are scarce (typically only a few in the protein history) but widespread (the majority of proteins experienced at least one). Typical coevolving residues are evolving slowly, are located in the protein core outside secondary structure motifs, and are more often in contact than expected by chance, even after accounting for their evolutionary rate and solvent exposure. An exception to this general scheme are residues coevolving for charge compensation, which are evolving faster than non-coevolving sites, in contradiction with predictions from simple coevolutionary models, but similar to stem pairs in RNA. While sites with a significant pattern of coevolution by compensatory mutations are rare, the comparative analysis of hundreds of structures ultimately permits a better understanding of the link between the three-dimensional structure of a protein and its fitness landscape.

## Introduction

The function of a biological molecule depends on its structure. It results that the structural characteristics of biomolecules impact the fitness effect of mutations in the genes that encode them and, therefore, determine their fate. The impact of structure on the process of molecular evolution has been extensively documented, both in RNA and proteins (Chen et al. 1999; Liberles et al. 2012; Moutinho et al. 2019). The structure and function of macromolecules, however, stem from the complex interactions – rather than the sum of the properties – of the underlying residues. As a consequence, the fitness effect of a mutation at a given position may depend on the state of the interacting residues, inducing a non-independent evolution, or coevolution (Starr and Thornton 2016). The interest of studying the signature of coevolution in molecular sequences has long been recognized, as detecting coevolving positions has the potential to point at functionally and structurally important interactions (de Juan et al. 2013).

Detecting coevolving positions is a statistically complex task, as the evolutionary process is not directly observable. Only its result is, in the form of extant sequences. The study of molecular coevolution is, therefore, grounded in phylogenetic comparative analysis: the shared history of species induces correlations in the sampled sequences that need to be disentangled from the functional correlations between sites. Furthermore, the geometry of interactions may vary in time and sequence space, not necessarily involving the same set of residues at different time points during the evolutionary history of the molecule. A large corpus of methods have been developed in order to address (some of) these issues, some using explicit evolutionary modelling of coevolution (e.g. (Pollock et al. 1999; Dib et al. 2014; Behdenna et al. 2016)), others relying on increasingly large data sets and advanced data mining procedures (Weigt et al. 2009; Jones et al. 2012; Wang et al. 2017; Li et al. 2019) to assess the patterns of site covariation. Furthermore, several case studies have provided a detailed understanding of the structural mechanisms of particular compensatory mutations (Ivankov et al. 2014; Storz 2018). The distribution of coevolving positions in proteins, however, and the underlying molecular mechanisms of coevolution remain largely unknown. A reason underlying this state-of-the-art is that model-based approaches are computationally intensive, preventing large scale comparisons, and typically produce small numbers of candidate coevolving positions with strong statistical support. Conversely, data mining approaches are able to predict the interaction network of residues in a molecule with good accuracy, providing that a very large number of sequences is available, thus preventing the use of such methods on a large variety of gene families and protein structures: only few gene families will match the necessary sample size. Furthermore, these methods are typically calibrated to detect physically interacting residues, which may or may not be coevolving, excluding potentially coevolving residues not in direct physical contact.

There is an apparent discrepancy between evolutionary methods, which predict relatively few coevolving positions, and pattern-based methods, which successfully predict a large proportion of residues that are physically in contact. The first point is explained by a theoretical argument (Ivankov et al. 2014; Talavera et al. 2015): for compensating mutations to occur, the first mutation must be sufficiently deleterious for the second mutation to be advantageous and invade the population. If the first mutation is too deleterious, however, it will be removed quickly from the population and will not be compensated. As a result, compensating mutations are predicted to be relatively rare, and coevolving sites should evolve rather slowly. Talavera and colleagues (Talavera et al. 2015) further argue that covariation methods primarily infer slowly evolving sites, which tend to be located at the core of proteins and are, therefore, more likely to be in close proximity. Paradoxically, in RNA molecules, interacting pairs within double-stranded helices have long been recognized as exhibiting a clear pattern of coevolution resulting from Watson-Crick interactions and show an accelerated rate of evolution compared to single-stranded regions (Smit et al. 2007). The frequency of compensating mutations in proteins, their distribution with regard to structural properties, and their relation to the evolutionary rate of coevolving sites remain to be established (Starr and Thornton 2016).

Here, we aim at generating an atlas of protein coevolution. We gathered a large dataset of sequence alignments of protein families for which at least one protein structure has been experimentally determined. We use a phylogenetic method that exhibits positions undergoing substitutions on the same branches of the phylogeny. (Dutheil et al. 2005; Dutheil and Galtier 2007). We further restrict our analysis to the case of coevolution by compensating mutations, where the first mutation at a given position has a negative fitness effect, for instance, because it results in a less stable protein structure, which is compensated by a second mutation at another position in the protein. In order to predict compensating mutations, we need a proxy for the fitness effect of single mutations. Considering that the fitness impact of a mutation depends on the biochemical properties of the encoded amino acids, we can predict this effect by quantifying the change in such properties (Grantham 1974). Neher (Neher 1994) first introduced this idea, developing a method looking at positive and negative correlations of biochemical properties between positions of sequence alignments. Dutheil and Galtier (Dutheil and Galtier 2007) introduced a measure of phylogenetic compensation (referred to as the “compensation index”), which assesses the conservation of a biochemical property in a group of sites, given their individual variation. More specifically, considering *n* sites and a phylogeny with *m* branches, we note 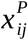 the change of a biochemical property *P* for site *i* on branch *j*. We further note 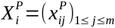 the vector of branch-specific change for site *i*. The compensation index *C*(*G*) for a group of sites *G* is then computed as

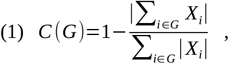

where |*X*| denotes the norm of *X* vector. When the changes at a given set of positions tend to be in opposite directions, |∑ *X*_*i*_| tends towards 0 and C towards 1. Conversely, if the changes are in the same direction, |∑ *X*_*i*_| equals ∑|*X*_*i*_| and C is equal to 0. The CoMap method uses a model-based substitution mapping procedure to infer the changes at each site on each branch of the phylogeny (Tataru and Hobolth 2011), combined with a clustering approach to detect candidate coevolving groups, which are then tested using simulations under the null hypothesis of independently evolving positions (Dutheil and Galtier 2007). While these simulations assume site independence, they conserve other aspects of the evolutionary process, such as the phylogenetic relationship of sequences, the probabilities of individual amino acid substitutions, and the site-specific rate of substitutions, allowing a precise evaluation of the coevolution signal while accounting for potentially confounding factors. Applying this method to hundreds of protein families and statistically analyzing hundreds of thousands of residues with structural annotations, we determine the drivers of protein intra-molecular coevolution.

## Results

We gathered a large data set of protein sequence families from complete genome sequences and for which structural information was available for at least one representative sequence. We initially analyzed gene families from the Archaea, Eukaryotes and Bacteria separately. However, we later restricted our analyses to bacterial families due to the small number of sequences and structures retained in the two other domains. We developed a stringent pipeline controlling for alignment uncertainty and sequence redundancy (see Material and Methods). Our curated data set contains 1,630 protein families, with a number of sequences per family ranging from 100 to 400 (Supplementary Table S1). In order to assess the compensating nature of substitutions, we considered several biochemical properties. Following previous studies, we considered the volume, polarity and charge of amino acid residues (Neher 1994; Pollock et al. 1999; Tufféry and Darlu 2000). The AAIndex database (Kawashima et al. 2008), however, contains more than 500 non-independent indices. Using clustering techniques, Saha et al. (Saha et al. 2012) have shown that these redundant properties are clustered into eight groups. We included the eight indices corresponding to the centres of these clusters (hereby referred to as “synthetic indices”), in order to provide an objective and comprehensive measure of amino acid biochemical properties. These indices are defined as follow (Saha et al. 2012): electric properties (I1), hydrophobicity (I2), alpha and turn propensities (I3), physicochemical properties (I4), residue propensity (I5), composition (I6), beta propensity (I7), and intrinsic propensities (I8). We ran the CoMap coevolution detection method to detect non-overlapping groups of coevolving positions, with a size ranging from 2 to 10, for the 11 biochemical properties. As a result, each site was annotated as coevolving for a given property if it belonged to a significant group. Structural properties, such as secondary structure motif and solvent exposure, as well as evolutionary rates, were recorded for each analyzed site (see Material and Methods).

### Substitution mapping enables the detection of compensating mutations

We detail two candidate groups to illustrate the nature of the positions predicted as coevolving in our data set. The first group involves a pair of distant sites at positions 146 and 237 in the sequence alignment of protein family HOG000218359. The three-dimensional structure used as a representative is the Menaquinone biosynthesis MenD protein of *Bacillus subtilis* (PDB ID: 2X7J). The protein is a tetramer (Dawson et al. 2010), and the A chain was used as a reference. The coevolving positions correspond to position Glu-109 and Arg-174 in the *B. subtilis* sequence (Figure 1A). The positions are significantly coevolving for charge compensation (P-value = 4.940333e-06, significant after correction for multiple testing with a false discovery rate (FDR) of 1%). While the two sites show low substitution rates, they undergo two significant changes in two branches of the tree (Figure 1B). These changes show a perfect compensation signal, from a positively charged residue to a negatively charged residue at one position and from a negatively charged residue to a positively charged residue at the other position (Figure 1B). This signal is illustrated on the “compensogram” in Figure 1D (see Material and Methods): the compensation is perfect (C = 1) for the first branch and almost perfect for the second branch (C > 0.9).

**Figure 1:**
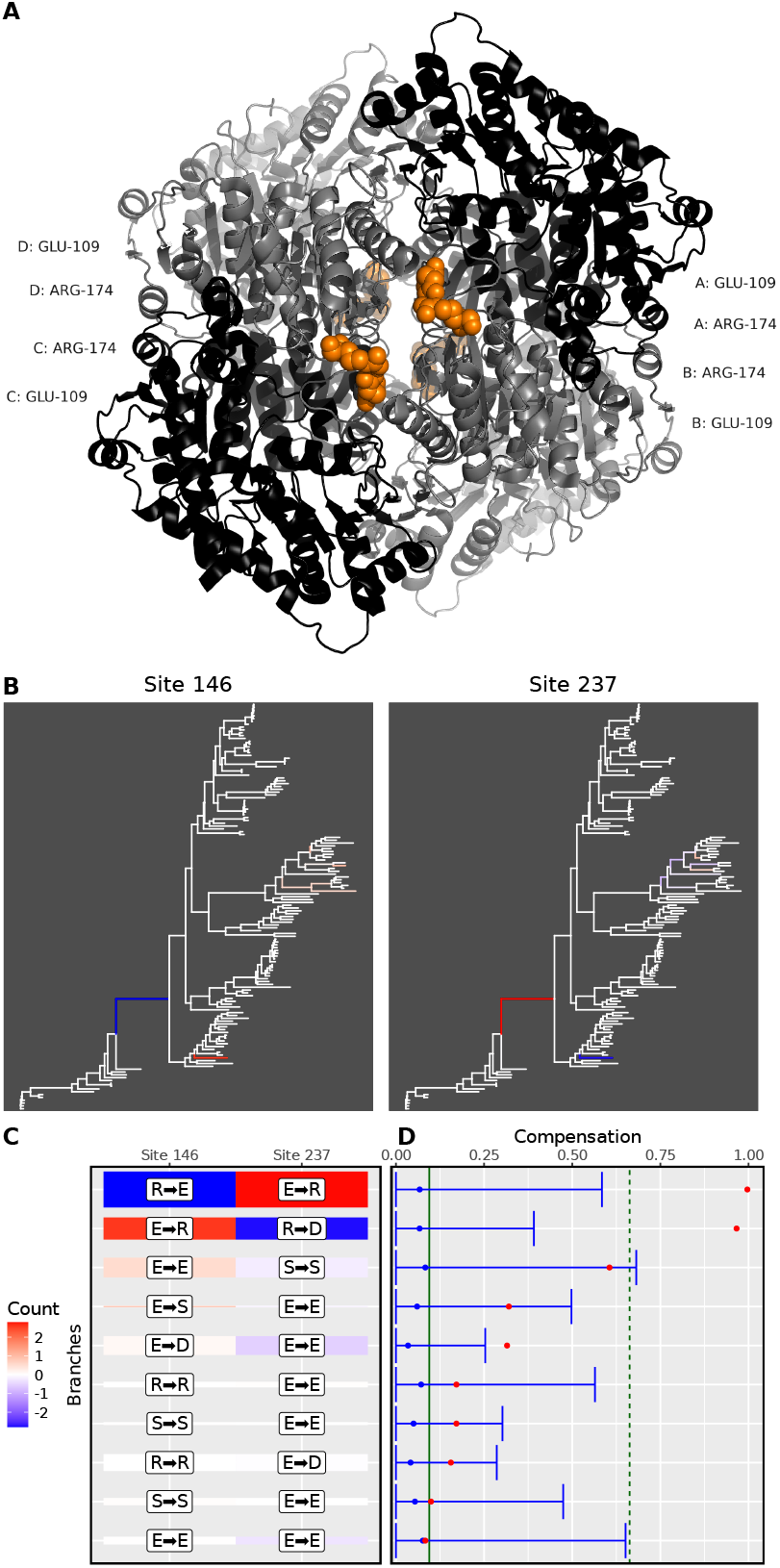
Example 1, positions Glu-109 and Arg-174 of the Menaquinone biosynthesis MenD protein. **A:** three-dimensional structure of the Menaquinone biosynthesis MenD tetramer. Residues at coevolving positions are shown in spacefill and colored in orange, for each monomer. **B:** estimated amounts of charge change are plotted on the branches of the phylogeny for the two sites. Negative changes (i.e., charge reduction) are plotted with a blue gradient, while positive changes (i.e., charge increase) are colored with a red gradient. **C:** Heatmap showing the changes for the top 10 branches, ranked according to the amount of charge compensation. Colors as in panel B. The height of the tile is proportional to the total branch length. The marginal maximum likelihood ancestral state reconstruction is indicated within white labels but was not used for the compensation calculations, as these were integrated over all possible ancestral states, weighted by their respective likelihood values. **D:** compensogram of the two sites for the top 10 branches, as defined in B. Red dots indicate the charge compensation index for each branch. Blue dots and error bars indicate the mean and 95% confidence interval of the site-permutation test, for each branch. The green vertical solid and dash lines represent the mean and 95% upper bound of the branch permutation test (see Material and Methods).

The second example group involves three positions from the HOG000227724 family, whose representative structure is the A-chain of the dTDP-6-deoxy-D-xylo-4-hexulose 3,5 epimerase (RmlC) of *Salmonella typhimurium* (PDB ID: 1DZR, Figure 2A) (Giraud et al. 2000). The coevolving positions include the two close residues Glu-16 and Phe-20, as well as Asn-150. The three residues are located at the surface of the protein but are not in contact. They were detected as coevolving for volume compensation (P-value = 0.0007818069), with several branches showing substitutions leading to a change in residue volume (Figure 2B). At least six branches show a significant signal of compensation (Figure 2C-D). The most significant branch show a mutation from soleucine (Grantham’s volume: 111, large) to alanine (Grantham’s volume: 31, small, -72% volume change) at site 63, compensated by two substitutions at site 59 and 208: glutamine (Grantham’s volume: 85, medium) to arginine (Grantham’s volume: 124, large, +46% volume change) and glutamic acid (Grantham’s volume: 83, medium) to isoleucine (Grantham’s volume: 111, large, +34% volume change). The total volume of the three sites changes from 279 to 266, which only represents a - 5% volume change, illustrating the compensatory nature of the substitutions and the high compensation index (C > 0.9). These two examples illustrate the substitution patterns of the sites detected as coevolving. In particular, the coevolution signal is here defined in a phylogenetic context and can be traced back to individual substitutions.

**Figure 2:**
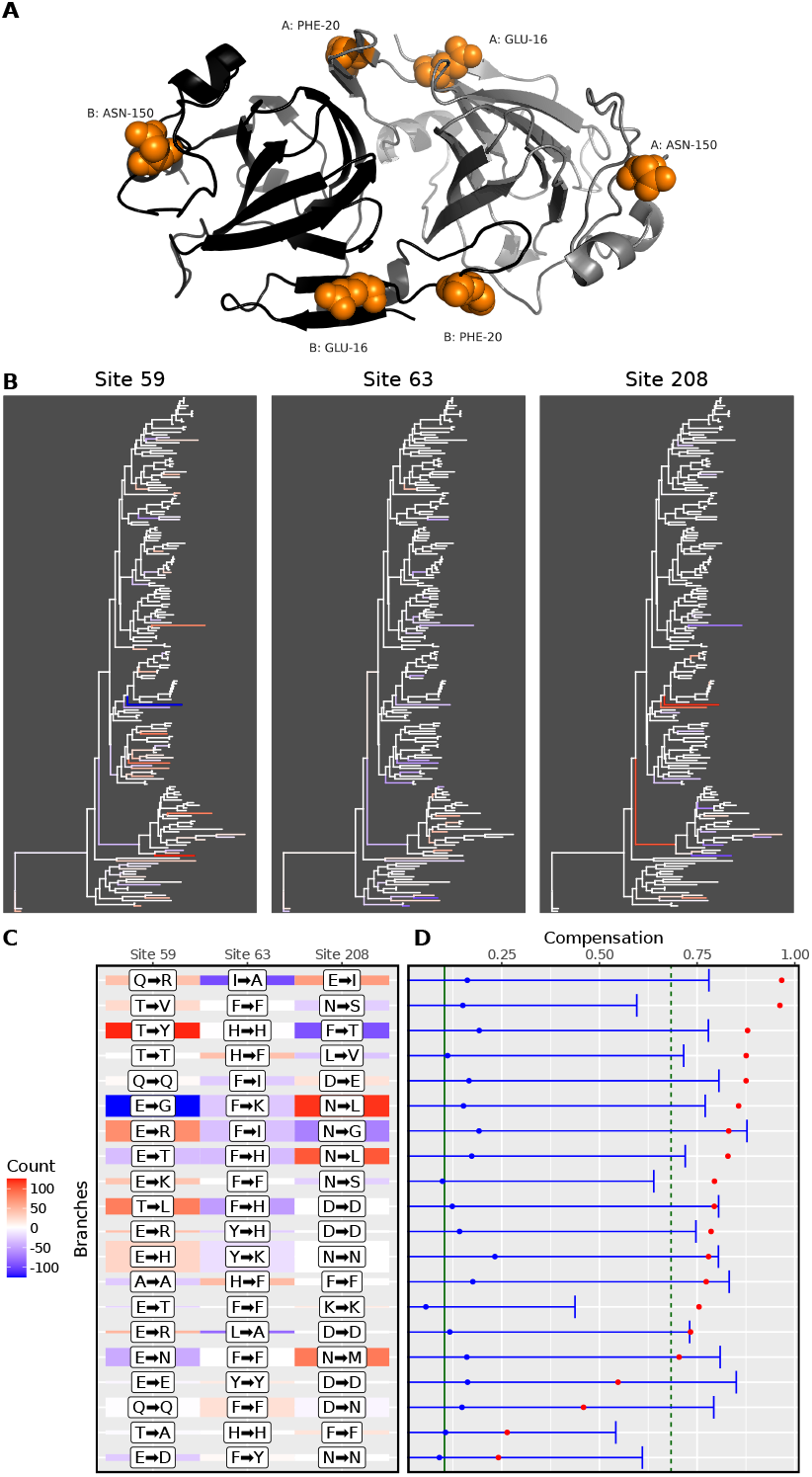
Example 2, positions Glu-16, Phe-20 and Asn-150 of the dTDP-6-deoxy-D-xylo-4-hexulose 3,5 epimerase (RmlC) protein. **A:** three-dimensional structure of RmlC dimer. Residues at coevolving positions are shown in full and colored in orange, for each monomer. **B:** changes of volume are plotted on the branches of the phylogeny for the three sites. Negative changes (i.e., volume reduction) are indicated in blue, while positive changes (i.e., volume increase) are colored in red. **C:** Heatmap showing the changes for the top 20 branches, ranked according to the amount of volume compensation. Colors as in panel B. The height of the tile is proportional to the total branch length. The marginal maximum likelihood ancestral state reconstruction is indicated within white labels but was not used for the compensation calculations, as these were integrated over all possible ancestral states, weighted by their respective likelihood values. **D:** compensogram of the two sites for the top 20 branches, as defined in B. Red dots indicate the charge compensation index for each branch. Blue dots and error bars indicate the mean and 95% confidence interval of the site-permutation test for each branch. The green vertical solid and dash lines represent the mean and 95% upper bound of the branch permutation test (see Material and Methods).

### Coevolution is scarce within, but widespread among protein families

Our coevolution analysis encompasses 366,794 sites, among which 51,661 (14%) were found to be coevolving for at least one biochemical property (Figure 3). While most sites were coevolving for a single biochemical property, the number of sites found to be coevolving by at least two methods was significantly greater than expected by chance (permutation test, P-value < 1e-4). This significant overlap may be explained by some sites being more likely to be detected as coevolving than others, either because of heterogeneous statistical power across sites, or their underlying functional and structural properties. We note that the eight synthetic indices detected fewer coevolving sites than the *classical* properties volume, polarity and charge. This result suggests that all biochemical properties are not as likely to inducing coevolution. Interestingly, the two properties for which a coevolutionary scenario is perhaps most intuitive (volume, big-to-small compensated by small-to-big mutations, and charge, positive-to-negative compensated by negative-to-positive mutations) were the ones leading to the largest number of detected positions. The coevolving sites predicted with these properties encompassed more than a third (34%) of all predicted positions.

**Figure 3:**
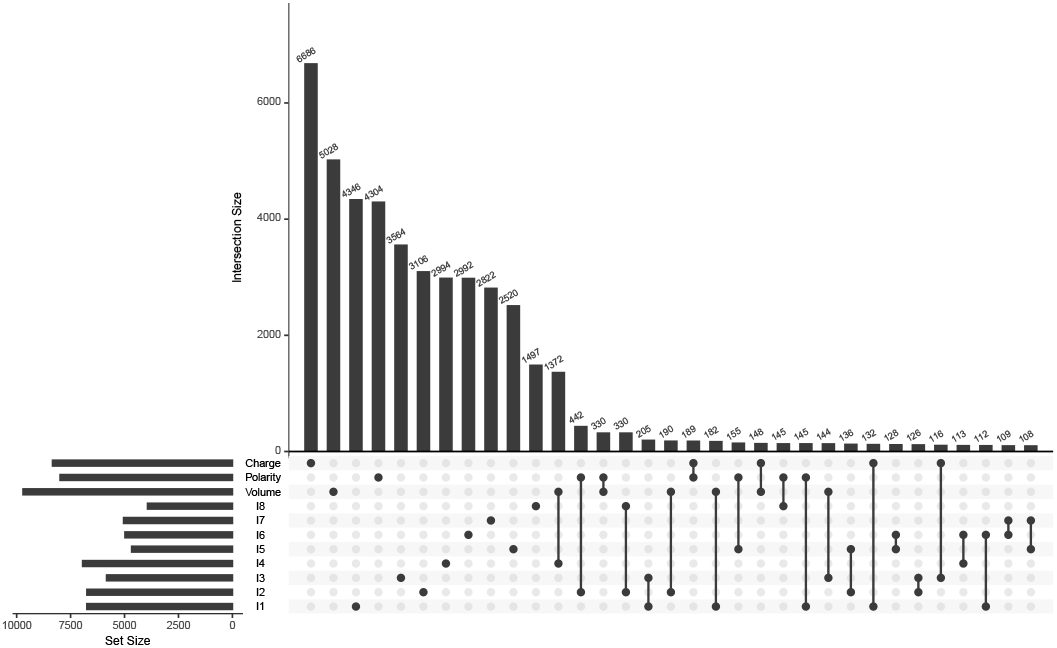
Upset diagram showing the overlap of sites detected as coevolving for compensation of various biochemical properties. The numbers of sites detected as coevolving (set size) are shown on the bottom left barplot. The numbers of sites detected by one or several methods (intersection size) are shown as the top barplot, sorted by decreasing size.

The number of detected coevolving positions in a protein family linearly increases with the size of the number of analyzable sites when this number is below ca 230 amino acids but decreases when the number of sites exceeds this threshold (Supplementary Figure S1A). Two opposite trends can explain this relation. Larger proteins have a larger number of residues interactions and offer more possibilities for coevolution. Conversely, they are also more conserved, meaning that mutations occurring in these proteins tend to be more deleterious and removed from the population, leaving less chance for compensating mutations to occur. In agreement with this hypothesis, we find a negative correlation between protein length and the tree diameter of each family (Kendall’s tau = -0.27, P-value < 2.2e-16, Supplementary Figure S1B), and a positive correlation between the tree diameter and the proportion of coevolving sites per family (Kendall’s tau = 0.17, P-value < 2.2e-16, Supplementary Figure S1C).

The evidence for coevolution within proteins was globally scarce since only 16% of the analyzed positions were involved in a coevolving group (median of all protein families). Coevolution, however, was found to be a general mechanism among proteins: 98% of the protein families that we analyzed had at least one coevolving group. In the following, we combined all positions from all protein families and unravelled the factors determining the occurrence of coevolution.

### Coevolving positions are evolving slowly unless they are coevolving because of charge compensation

We assessed the evolutionary rate of coevolving positions and tested whether the detected positions were more conserved, as predicted by the coevolution model of Talavera et al. (Talavera et al. 2015). An alternative hypothesis is that fast-evolving positions are more likely to be detected as coevolving because of the increased statistical power stemming from the larger number of underlying mutations (Dutheil 2012). For each biochemical property, we compared the evolutionary rate (see Material and Methods) for sites predicted to be coevolving or not. For all properties, coevolving positions had a lower evolutionary rate, except the charge property for which coevolving sites evolved faster than non-coevolving sites (Figure 4). A particularity of the charge property is its discrete nature: each amino acid is characterized by one of three possible states – positively charged, negatively charged, or neutral – while other properties are continuous. To test whether this difference in property nature could affect the capacity to detect coevolving sites, we discretized the volume and polarity indices in three categories (see Material and Methods) and rerun the coevolution detection analysis. We found that positions coevolving for volume and polarity had a lower rate of evolution, even when considering only three categories, suggesting that the discrete nature of the property is not responsible for the inferred faster evolution of coevolving charged residues (Supplementary Figure S2). We next aimed at assessing which structural characteristic impacts the propensity to coevolve, accounting for the underlying evolutionary rate of the sites.

**Figure 4:**
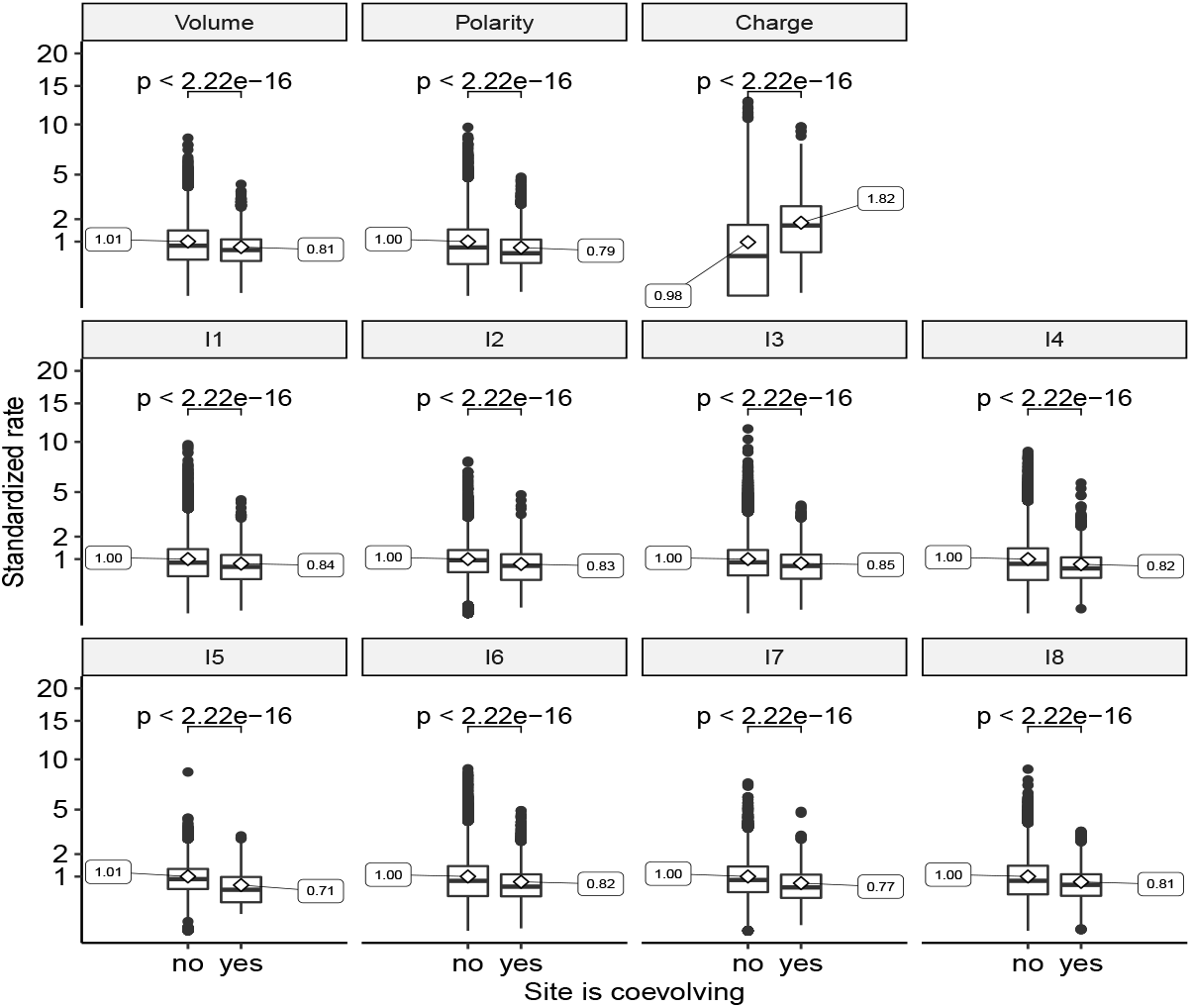
Evolutionary rates of coevolving sites. Box-whiskers plots (first and third quartile, medians) of standardized rates for coevolving vs non-coevolving sites for each biochemical property. Mean values of each distribution are indicated as side labels. P-values of corresponding Wilcoxon rank tests are indicated on top of each comparison.

### There is comparatively less coevolution within secondary structure motifs

We used mixed generalized linear models to assess the impact of structural properties on the propensity of sites to coevolve. Each biochemical property was analyzed separately, and each site was considered an independent data point. The protein family was treated as a random effect, allowing sites within the same protein to share the same error distribution, while different protein families may have distinct ones. Whether each site was found to be coevolving was used as a binary response variable. Structural properties of each site were recorded, including its relative solvent accessibility (RSA), its secondary structure motif (one of α-helix, 3-10 helix, π-helix, strand, β-bridge, turn, bend, intrinsically disordered or unknown), and the evolutionary rate of the site. Whilst several explanatory variables are potentially intrinsically correlated, all variance inflation factors were found to be close to 1.0 and much lower than 5, an empirical threshold for identifying colinearity issues (James et al. 2013). The highest values were observed for the RSA variable, for which they are above 2.0 (Supplementary Table S2).

The evolutionary rate of the sites was found to be the most significant variable (Table 1). Consistent with the results above, the effect was significantly negative for all properties except charge, where it was significantly positive. Solvent exposure was generally found to have a negative impact on coevolution, meaning that coevolving residues tend to be buried. Indices 1 and 3 are an exception, though, as exposed residues have an increased probability of experiencing mutations that compensate for electric properties or alpha and turn propensities. Conversely, secondary structure was found to have little impact on the occurrence of compensating mutations. The most substantial effect was observed for β-strands and α-helices, where it was consistently negative: sites in helices and strands are less likely to experience compensating mutations. We further note a significant positive interaction between solvent exposure and strand for properties “polarity”, “charge”, and index 1, indicating that for these amino acid properties, exposed residues in strands have an increased probability of coevolving. The property for which secondary structure has the strongest effect was index 3, “alpha and turn propensities”, showing a significant positive effect in turn and bend motifs, as well as in disordered regions. This index reflects the propensity of amino acids to be found in helices (high index values) vs turns (low index values). Bends, turns and disordered regions prefer amino acids with low values for index 3. A potential coevolution scenario would involve that a mutation of a preferred amino acid type toward a non-preferred type may be compensated by a mutation at another position within the same region, involving a non-preferred type toward a preferred amino acid type. We also report a significant negative interaction with solvent exposure in these three types of regions, suggesting that the chances for coevolution are higher for buried residues.

**Table 1:** Impact of structural properties on the probability of sites to coevolve. For each biochemical property, a generalized linear model with mixed effects was fitted. Whether a site was found coevolving according to the biochemical property was set as a binary response variable, and the evolutionary rate and structural properties were set as explanatory variables. The protein family was set as a random factor.

|  | variable | volume | polarity | charge | I1 | I2 | I3 | I4 | I5 | I6 | I7 | I8 |
| --- | --- | --- | --- | --- | --- | --- | --- | --- | --- | --- | --- | --- |
|  | RE(Intercept) | 1.17 (***) | 1.23 (***) | 1.36 (***) | 1.68 (***) | 1.51 (***) | 1.69 (***) | 1.82 (***) | 2.18 (***) | 2.09 (***) | 2.34 (***) | 2.58 (***) |
|  | (Intercept) | -3.45 (***) | -3.8 (***) | -4.94 (***) | -3.63 (***) | -4.09 (***) | -4.24 (***) | -4.25 (***) | -5.18 (***) | -4.74 (***) | -5.05 (***) | -5.67 (***) |
|  | Np<sup>1</sup> | -0.35 (***) | -0.38 (***) | 0.52 (***) | -1.44 (***) | -0.34 (***) | -0.71 (***) | -0.34 (***) | -0.34 (***) | -0.61 (***) | -0.34 (***) | -0.31 (***) |
|  | Rsa<sup>2</sup> | -0.7 (***) | -0.33 (*) | -0.32 (*) | 0.4 (**) | -0.34 (*) | 0.34 (*) | -0.51 (***) | -0.28 | -0.54 (**) | -0.69 (***) | -0.44 (*) |
| Secondary structure | α-helix | -0.17 (***) | -0.17 (**) | 0.01 | -0.11 (*) | -0.15 (**) | -0.17 (**) | -0.08 | 0 | 0 | -0.24 (***) | -0.24 (**) |
|  | 3-10 helix | 0.17 (*) | 0.24 (*) | -0.03 | 0.19 (.) | -0.09 | 0.2 (.) | 0.07 | 0.13 | 0.18 | -0.2 | -0.2 |
|  | π-helix | -0.02 | 0.26 | 0.25 | -0.3 | 0.24 | 0.52 (*) | 0.14 | 0.48 (.) | 0.04 | 0.09 | 0.05 |
|  | Strand | -0.21 (***) | -0.26 (***) | -0.31 (***) | -0.17 (**) | -0.17 (**) | -0.26 (***) | -0.13 (*) | -0.05 | -0.09 | -0.11 | -0.18 (*) |
|  | β-bridge | -0.24 | -0.23 | -0.15 | 0.03 | -0.09 | -0.13 | -0.23 | 0 | -0.04 | 0.04 | -0.07 |
|  | Turn | 0.16 (*) | 0.14 (.) | 0.16 (.) | 0.06 | -0.06 | 0.18 (*) | 0.2 (*) | -0.03 | 0.12 | -0.09 | -0.06 |
|  | Bend | 0.12 (.) | 0.11 | 0.24 (*) | 0.08 | -0.06 | 0.24 (**) | 0.14 (.) | 0.14 | 0.04 | -0.23 (*) | -0.03 |
|  | Disordered | -0.22 (*) | 0.09 | 0.25 (*) | 0.12 | -0.08 | 0.4 (***) | -0.02 | -0.03 | 0.06 | -0.13 | -0.11 |
| Rsa:secondary structure | α-helix | -0.22 | 0.11 | 0.14 | 0.28 | -0.17 | -0.45 (*) | -0.52 (**) | -0.38 (.) | 0.15 | 0.01 | -0.17 |
|  | 3-10 helix | -0.37 | -0.54 (.) | 0.1 | -0.2 | 0.2 | -0.71 (*) | -0.32 | -0.08 | 0.24 | 0.74 (*) | 0.45 |
|  | π-helix | -0.29 | -0.16 | 0.15 | 1.77 (*) | -0.2 | -2.19 (.) | -0.59 | -1.26 | 0.41 | 0.43 | 0.48 |
|  | Strand | 0.13 | 0.68 (***) | 1.22 (***) | 0.8 (***) | 0.29 | 0.06 | -0.06 | 0.37 | 0.38 | 0.35 | 0.23 |
|  | β-bridge | 0.47 | 0.91 | 0.03 | 0.86 | -0.08 | 0.39 | -0.51 | 0.39 | -0.46 | -0.22 | 0.25 |
|  | Turn | -0.23 | -0.53 (*) | -0.39 (.) | -0.44 (*) | -0.35 | -0.65 (**) | -0.53 (*) | -0.19 | -0.12 | -0.06 | -0.15 |
|  | Bend | -0.05 | -0.14 | -0.31 | -0.09 | 0.11 | -0.72 (**) | -0.29 | -0.39 | 0.07 | 0.24 | -0.13 |
|  | Disordered | 0.59 (*) | -0.11 | -0.23 | -0.12 | 0.13 | -0.65 (*) | 0.04 | 0.11 | 0.08 | 0.52 (.) | 0.03 |

<sup>1</sup> Standardized norm of the weighted substitution vector (evolutionary rate)
<sup>2</sup> Relative solvent accessibility (solvent exposure)

In the following, we more specifically studied the role of secondary and tertiary structure in the occurrence of compensatory mutations. First, we assessed whether the proximity of residues in the three-dimensional structure impacted the occurrence of coevolution. Second, we tested whether coevolving residues located in a secondary structure motif tended to be located in the same element (helix, strand, or sheet). To test these hypotheses, we developed a permutation test controlling for evolutionary rate and solvent exposure.

### Coevolving residues are more likely to be in contact, independently of their exposure or evolutionary rate

Residues at the core of proteins (low RSA) tend to be more conserved and more connected (Supplementary Figure S3). In order to test whether coevolving residues are more often in contact than randomly selected sites, we need to compare positions with similar rates or solvent exposure. For that purpose, we developed a Monte-Carlo algorithm that samples groups of sites conditioned on a third variable (see Material and Methods). As expected, we showed that sampling residues without conditioning led to a bias: randomly selected sites had a higher rate and exposure on average than the groups of sites detected as coevolving by all methods but the one accounting for the charge property, for which the opposite pattern was observed. This bias is successfully removed by using conditional sampling (Supplementary Figure S4). These results align with the observation that coevolving sites evolve more slowly and are buried, while sites detected as coevolving for charge evolve faster and are exposed.

Using the conditional sampling algorithm, we tested whether sites detected as coevolving were closer to each other in the three-dimensional structure than random sites. We computed the average pairwise three-dimensional distance between the alpha-carbons of the residues within each detected group, which we averaged over all groups. We then computed the same statistic over 1,000 random sets of groups with the same sizes and similar rates or RSA. Only 0.35% of all site groups were excluded from the randomization test due to the lack of sites with a similar rate within the same protein. Conversely, 31% had to be discarded when conditioning on RSA. We found that the observed statistic is significantly lower than in the case of random sets for all methods (Figure 5A, all P-values < 0.001), suggesting that coevolving residues are in closer proximity than expected by chance. To further assess whether this proximity is explained by residues being in physical contact, we investigated the connectivity graph of the residues in each group. As an approximation, we considered two residues to be in contact if their α-carbon distance was shorter than 8 Å and computed the proximity graph of the residues, where an edge connects two residues if they are in contact. To measure the number of contacts, we counted the number of graphs for the group: this number is equal to 1 if all residues are in contact, either directly or indirectly. If no residue is in contact with any other, the number of graphs equals the number of residues. We further standardized this measure so that it is comprised between 0 and 1 to be comparable between groups of different sizes (see Material and Methods). We showed that this statistic is lower than expected by chance when sampling groups of sites with a similar rate or RSA (Figure 5B, all P-values < 0.001). Therefore, we conclude that coevolving residues are more likely to be in contact in the three-dimensional structure. This effect is not due to their slow evolutionary rate and low solvent exposure.

**Figure 5:**
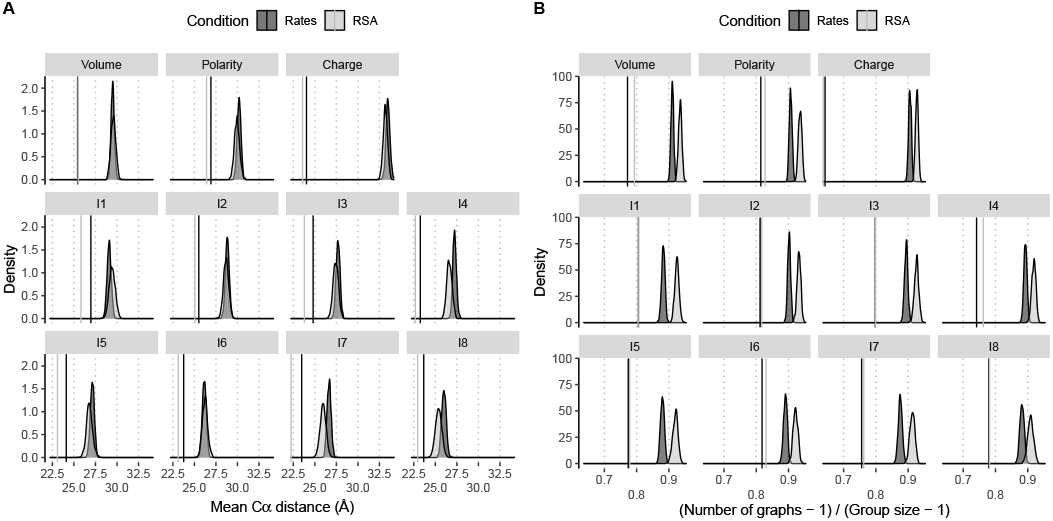
Three-dimensional proximity of coevolving sites. Average Ca distance (A) and relative number of contact groups (B) of all groups detected to be coevolving by compensation for each biochemical property. Densities are computed from 1,000 samplings, conditioned over the evolutionary rate or the solvent exposure (see Material and Methods). Only groups for which sites with a similar rate or RSA values could be sampled were included. Observed values are indicated as vertical bars and are all significantly lower than expected by chance, with a P-value < 1/1001.

Using a similar approach, we further investigated whether coevolving residues in a given secondary structure motif were located in the same element. We generated two sub-datasets, only containing sites in helices or strands, respectively. We then computed the connectivity graphs in each case and considered two residues in contact if they were in the same helix, strand, or sheet. By sampling among helix or strand sites within each protein family, we computed the expected distribution of these statistics. We also compared the results with randomization where sites were constrained to have similar evolutionary rate or RSA. We report consistent results between these analyses (Figure 6): coevolving sites located within helices are significantly more often in the same helix than expected by chance (Figure 6A).

**Figure 6:**
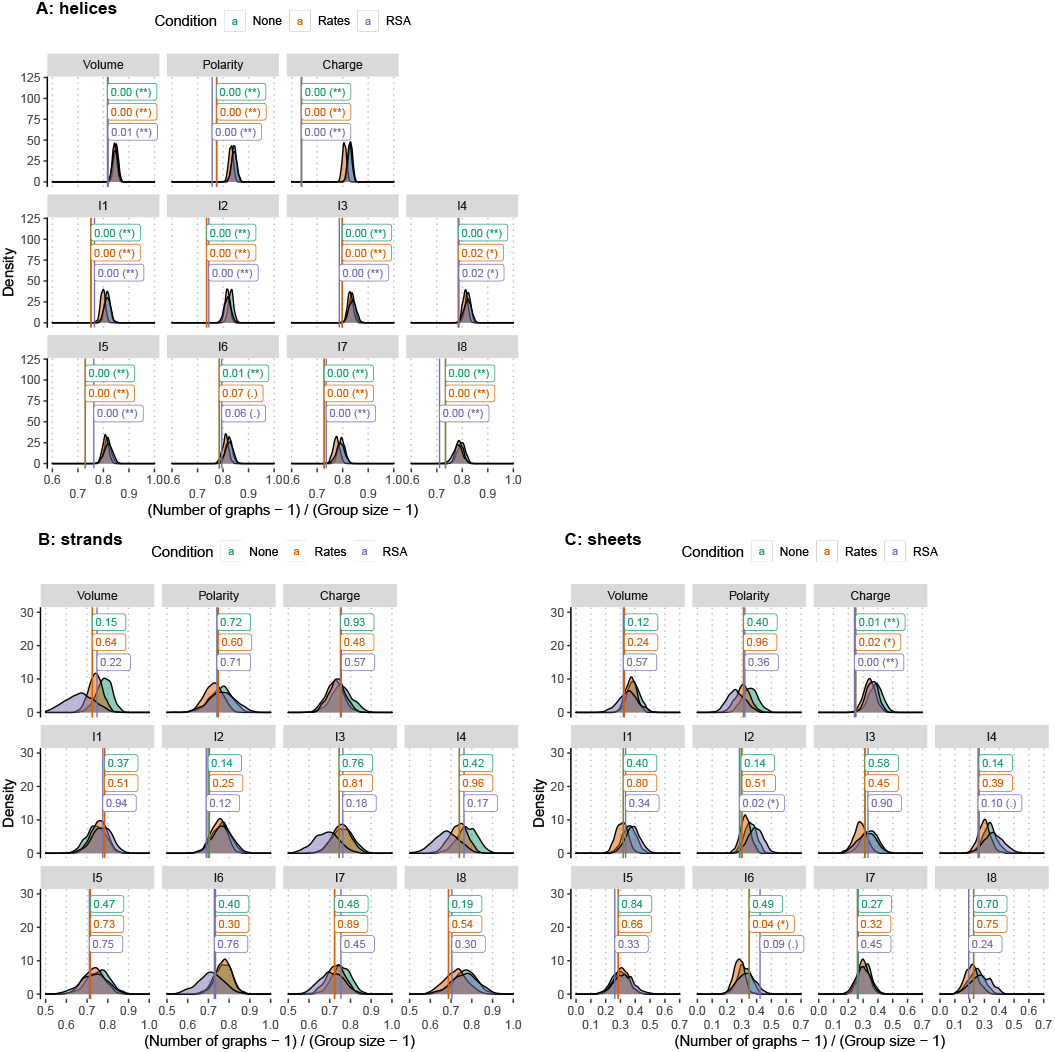
Are coevolving positions located within the same secondary structure element? A) Helices, B) strands, and C) sheets. The x-axis measures the relative number of secondary structure elements: a value of 0 indicates that all sites within a coevolving group are in the same element, while a value of 1 is obtained when each site is in a distinct element. Densities are computed from 1,000 samplings, conditioned over the secondary structure element alone (None), secondary structure and evolutionary rate (Rate), or secondary structure and solvent exposure (RSA, see Material and Methods). Only groups for which sites with a similar rate or RSA values could be sampled were included. Observed values are indicated as vertical bars together with the corresponding P-values.

Conversely, coevolving sites within strands are not more often in the same one (Figure 6B). They are also generally not more often in the same sheet, except sites coevolving for charge (Figure 6C). These results allow us to refine our previous conclusions: while there are fewer compensating mutations in secondary structure motifs, this level of organization creates evolutionary constraints susceptible to induce coevolution. This is particularly clear for helices, while only charge compensation could be evidenced in the case of β-sheets.

## Discussion

We generated an atlas of protein compensating mutations, applying a phylogenetic method on a curated set of protein alignments for which a representative three-dimensional structure was available. We employed a substitution mapping procedure, inferring the position of all amino acid substitutions that occurred at every site of the alignment and every branch of the underlying phylogeny. We used indices of amino acid biochemical properties to build a proxy for the fitness effect of amino acid substitutions and assess their compensatory nature (Dutheil and Galtier 2007). The statistical procedure used to assess the significance of the coevolution signal accounts for the phylogenetic correlation and the site-specific rate of evolution, addressing some of the common pitfalls in predicting site associations (de Juan et al. 2013).

We found the number of positions with a significant coevolution signal to be generally low, and coevolving positions were generally evolving slowly, showing a higher-than-average level of conservation. These results indicate that compensating mutations are rare, as predicted by theoretical models (Talavera et al. 2015). Most of the families that we tested, however, exhibited some coevolving positions, showing that while rare, compensating mutations are widespread among proteins with distinct structural and functional properties. From a genetic perspective, the mechanism of compensating mutations implies that the negative fitness effect of a first mutation is compensated by the positive fitness effect of a second mutation. The fitness effect of the two mutations, therefore, depends on the order in which they occur: the same mutation may have a negative effect if it appears first or positive if it happens second, compensating for a preceding deleterious mutation. Compensating mutations constitute a particular case of non-additive fitness, termed reciprocal sign epistasis, which induces valleys of low fitness in fitness landscapes (see (Whitlock et al. 1995) for a review, and (Poelwijk et al. 2011)). Our empirical results suggest that reciprocal sign epistasis is a general feature of protein-coding gene fitness landscapes, independent of the function of the encoded proteins.

Our analyses confirm a strong link between protein structure and coevolution of residues (de Juan et al. 2013). In particular, our results highlight the importance of tertiary structure in shaping the fitness landscape of proteins: coevolving residues tend to be located outside secondary structure motifs, at the hydrophobic core of proteins, and generally in close proximity. Importantly, we show that the link between structural properties and patterns of sites coevolution is not a by-product of the structural constraints shaping site-specific evolutionary rates (Talavera et al. 2015): coevolving residues are more likely to be physically in contact than random sites with an equal rate or solvent exposure. While failing to account for evolutionary rates may lead to sites being falsely labelled as coevolving, in particular when large amounts of such sites are predicted for a given protein, this fact does not imply that compensating mutations do not occasionally occur and cannot be detected.

The patterns of coevolving positions with respect to site properties appeared to be consistent across the range of biochemical properties that we used as a fitness proxy. This robustness extends to the properties selected from a multi-dimensional analysis of more than 500 properties available, ensuring that our conclusions are not biased toward any prior assumption on which biochemical property might be relevant. A notable exception is the pattern of sites detected as coevolving for charge compensation, which have a higher substitution rate than non-coevolving sites. Coevolution for charge compensation mirrors the pattern of coevolution in RNA, where Watson-Crick pairs in double-stranded helices evolve at a faster rate than single-stranded regions and pairs interacting via non-Watson-Crick interactions (Dutheil et al. 2010). It is particularly intriguing that distinct biochemical constraints lead to opposite patterns of evolutionary rates, suggesting that the frequency at which fitness valleys are crossed in a fitness landscape cannot be fully predicted by simple models with fixed selection coefficients. A key to the understanding of these dynamics may lie in the accounting of selective constraints acting on intermediate organizational levels. This seems to be the case in RNA, where selective force may act at the level of double-stranded stems (Dutheil et al. 2010). Here, however, we show that secondary structure motifs comparatively seem to play little role in the occurrence of coevolving positions in proteins, where intermediate levels of organization pertinent to molecular coevolution are only starting to be characterized (Halabi et al. 2009).

## Material and Methods

### Sequence retrieval

Protein sequences were retrieved from the HOGENOM database (release 06) (Penel et al. 2009), which contains families of homologous protein-coding genes from completely sequenced genomes of Bacteria, Archaea and Eukaryotes. We sampled the sequences in all three domains of life and found that only Bacteria had enough sequences to provide sufficient signal for coevolution detection. Thus, we targeted members of the Bacteria domain for this study. Using the ‘query_win’ retrieval program (Gouy and Delmotte 2008), 2,047 families were selected for which at least one bacterial sequence had at least one experimentally solved structure available. All the Archaea and Eukaryotic sequences were removed from the selected families. For each family, species having more than one sequence were discarded to avoid the comparison of paralogous sequences. To ensure the comparison of families with similar statistical power, we discarded families with less than 100 sequences.

We developed a pipeline to process each protein family, with the goal to minimize the amount of missing data and maximize the alignment reliability:

1. **Approximate alignment and phylogeny:** a fast multiple sequence alignment for each family was performed using Clustal Omega (Sievers et al. 2011). For each alignment, a first phylogenetic tree was built with FastTree (Price et al. 2010).
2. **Minimization of missing data:** the phylogenetic tree obtained at step 1 was used to remove sequences from the alignment in order to maximize the number of sites with sufficient coverage using the ‘bppAlnOptim’ program (Dutheil and Figuet 2015). Unresolved characters were considered together with gaps in coverage calculations, and only sites without gaps were kept.
3. **High-quality alignment:** filtered families were realigned using both Muscle (Edgar 2004) and Clustal Omega (Sievers et al. 2011). We built a consensus alignment by computing sum-of-pairs scores (SPS) for each alignment site and masking sites with an SPS lower than 80%. These high-quality consensus alignments were used in the following analyses.
4. **Phylogenetic sampling:** we performed a two-step phylogenetic sampling, first to remove highly similar sequences, which do not carry a biological signal, and second to limit the maximum number of sequences in each family in order to reduce computational time. The ‘bppPhySamp” program from the Bio++ program suite (Dutheil and Boussau 2008) was used to ensure that sequences in each alignment were at least 1% different from the other sequences and that there is a maximum of 400 sequences per alignment. Families with less than 100 sequences after sampling were further discarded, leaving a total of 1,684 families.
5. **Accurate phylogeny reconstruction:** the resulting masked alignments were used to build a maximum likelihood phylogenetic tree, using the PhyML program (Guindon and Gascuel 2003) (Guindon et al. 2010), using the LG protein substitution model (Le and Gascuel 2008) and a gamma distribution with four rate classes (Yang 1994). The best tree from Nearest Neighbour Interchange and Subtree Pruning Regrafting algorithms (Guindon et al. 2010) was kept. The inferred phylogenies were used for the identification of coevolving sites.

### Identification of coevolving sites

Coevolving positions within each protein family was predicted using the CoMap package v1.5.2 (Dutheil and Galtier 2007). A false discovery rate (FDR) utilizing the built-in algorithm was then used to correct for multiple testing. In this study, 10,000 simulations per tested group were performed, and the groups with FDR < 1% were considered as coevolving. Compensation was used as a measure of coevolution which assesses the compensatory nature of cosubstitutions based on a given physicochemical property. The compensation indexes used as weights were retrieved from the AAindex database (Kawashima et al. 2008). Three commonly studied properties: Volume (as defined by Grantham, AAindex id: GRAR740103); Polarity (as defined by Grantham, AAindex id: GRAR740102); Charge (as defined by Klein, AAindex id: KLEP840101) were tested. In addition, we also tested eight non-overlapping properties (indexes) as described by Saha et al. (Saha et al. 2012). Saha et al. have categorized all published 544 amino acid indexes of the AAindex database into eight clusters, and these eights categories were used in this study. Corresponding categories are Electric property; Hydrophobicity; Alpha and turn propensities; Physicochemical properties; Residue propensity; Composition; Beta propensity and Intrinsic propensities. Because the charge property has three discrete states (positively charged, negatively charged and neutral), we further assess the impact of discrete vs continuous properties on the detection of coevolution by discretizing the volume and polarity properties. Residues G (volume (Grantham 1974): 3), A (31), S (32), and P (32.5) were considered as small, residues D (54), C (55), N (56), T (61), E (83), Q (85), and V (84) as medium, and residues H (96), M (105), I (111), L (111), K (119), R (124), F (132), Y (136), and W (170) as large. Residues L (polarity (Grantham 1974): 4.9), I (5.2), F (5.2), W (5.4), C (5.5), M (5.7), V (5.9), and Y (6.2) were classified as hydrophobic, residues P (8.0), A (8.1), T (8.6), G (9.0), and S (9.2) as intermediate, and residues H (10.4), Q (10.5), R (10.5), K (11.3), N (11.6), E (12.3), and D (13.0) as polar.

Out of 1,684 families, 24 families were discarded because likelihoods could not be computed because of numerical underflow. Numerical saturation, unavailability of secondary structure or solvent accessibility information (see below) resulted in a final total of 1,630 analyzable protein families (Supplementary Table S1). The diameter of each phylogenetic tree (maximum distance between any two leaves) as well as the median of sequence lengths for each family were computed using the ‘ape’ (Paradis et al. 2004) and ‘seqinr’ (Charif et al. 2005) packages for R, and recorded. The relationship between the number of detected sites and the alignment length was assessed using a broken regression model, as implemented in the ‘lm.br’ package for R (Adams 2017).

### Post-hoc tests of branch-specific compensatory substitutions

We developed two statistical post-hoc tests in order to better characterize the signal of coevolution for a group of candidate coevolving sites. Following the notations from the introduction, these tests compute the compensation statistic for a group of sites *G*, at a specific branch *j*. We note as 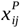 the change of a biochemical property *P* for site *i* on branch *j*. The compensation index *C*_*j*_(*G*) for a group of sites *G* on branch *j* is then computed as

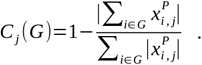

This measure is identical to the compensation index defined in the introduction for the full tree but applied to a single branch. We introduce two permutation tests to assess the significance of the compensation statistics. The first test randomizes the branches for each site, conditioning on the total amount of change at each site. The second test randomizes the sites for each branch, conditioning on the branch length. In both tests, the conservation statistic is computed for each permutation and its distribution compared to the observed value.

### Evolutionary rates

We measured the rate of molecular evolution according to a given biochemical property using the norm of the weighted substitution vector for each site (Dutheil and Galtier 2007). For a given site in the alignment, the number of substitutions that occurred on each branch of the phylogenetic tree was inferred by probabilistic substitution mapping. Each substitution type from an amino acid A into an amino acid B was then weighted by the absolute difference in the amino acid property for A and B.

### Structural information

The PDB ids of the structures associated with each family were extracted from the HOGENOM database and the corresponding PDB files retrieved from the Protein Data Bank (PDB) (Berman et al. 2000). For each alignment, all pairwise alignments of each homologous sequence with each subunit of each matching PDB structure were generated, and the PDB subunit leading to the highest alignment score was used as a reference structure for the protein family. The corresponding alignment was used to translate alignment positions into positions in the PDB structure. All scripts used for analyses were written in Python using the Biopython modules (Hamelryck and Manderick 2003). The reference PDB structures were then subsequently used to extract structural properties: secondary structure and relative solvent accessibility were obtained using the DSSP program (Kabsch and Sander 1983), run via the Biopython DSSP module. The distance of each residue to the solvent-accessible surface was computed via the MSMS program (Sanner et al. 1996), accessed via the ‘Biopython.ResidueDepth’ module. Both the residue depth and the C*a* depths were computed. We further computed the number of residues in contact with each residue using C*a* distance thresholds of 5, 8 and 10 Å. Because several of these properties may be intrinsically correlated, we examined their relationships using principal component analysis and non-parametric pairwise correlation (Spearman rank correlation). We found that the structural variables were highly correlated (Supplementary Figure S3): residue depths and numbers of contacts were highly positively correlated while highly negatively correlated with RSA. In summary, the more exposed a residue is, the closer it is to the surface of the protein and the less contacts it has with other residues. Because all these variables carry the same biological information, we only used RSA in the statistical analyses.

Intrinsic disorder measures for all residues were obtained using the DisEMBL version 1.4 program (Linding et al. 2003), after porting the original Python script to Python 3.0. We considered a site to be in a disordered region if its hotloop index was greater than 0.1204 and the site was not predicted by DSSP to be in a secondary structure motif. We considered sites labelled as disordered as not having any secondary structure, and we combined the results of DSSP and DisEMBL into a single discrete variable with states “no structure”, “alpha helix”, “3-10 helix”, “pi helix”, “strand”, “beta bridge”, “turn”, “bend”, and “disordered”.

In order to test whether the coevolving sites of a group are in proximity in the three-dimensional structure, we measured the distance between the two alpha carbon atoms of the residues. For groups with more than two sites, the average pairwise distance was computed. As an alternative measure of proximity, we considered residues with a distance ≤ 8 Å to be in contact. We then computed the number of subgroups of sites in contact within a group: if all residues are in contact with each other, then the number of subgroups is one. If each residue is distant from all others, then the number of subgroups is equal to the size of the group. We further standardized this measure by removing 1 and further dividing by the group size – 1, so that it is comprised between 0 (all residues in contact) to 1 (all residues apart). We note *N*_sub_ the resulting statistic, averaged over all candidate groups. Testing of residues proximity was performed using a dedicated Monte Carlo procedure. For each detected group in each protein family, a group of identical size was sampled among analyzed positions of the same protein family. In order to further enforce that sampled positions have evolutionary rates similar to the ones of the observed group, we performed a conditional sampling. For each site in each group, called the “focus” site, we sampled among the subset of sites with a rate no more than *x*% different for the rate of the focus site, *x* being an adjustable similarity threshold. We further ensured that sites were not sampled more than one time in each group, although a site in a protein was allowed to be sampled multiple times between distinct groups. Because the distribution of rates is typically skewed, we further restricted the sampled set of sites so that it contained as many values below and above the rate of the focus site. When less than five sites with similar rates were available, the group was not further considered in the randomization test. We first generated 100 replicates without conditioning, with a 10% similarity threshold and with a 20% threshold for comparison. We found that the 10% threshold removed any significant effect of the evolutionary rate (Supplementary Figure S4). We then generated 1,000 replicates per predicted group with the 10% threshold and computed p-values as (N+1)/1001, where N is the number of replicates with a measure greater or equal (or lower or equal, depending on the test) to the observed value in the data. A similar procedure was conducted using RSA instead of the evolutionary rate.

To test whether coevolving residues were located within the same secondary motif (helix, strand or sheet), we applied a similar randomization procedure after discarding all sites not located within secondary structures. In order to annotate sites with their secondary structure labels (helix and strand numbers), mmCIF files were used, and annotations parsed using scripts developed with the ‘BioPython.PDB’ package (Hamelryck and Manderick 2003). We computed the *N*_sub_ statistic (see above), considering whether sites are in the same motif or not. Sites in helices and strands were analyzed separately, and sampling was done in each case without further conditioning or by conditioning on evolutionary rates or RSA, with a 10% similarity threshold and with 1,000 replicates.

### Statistical analyses

All statistical analyses were conducted with the R statistical software (R Core Team 2020) using the ‘ggplot2’ (Wickham 2016), ‘cowplot’ (Wilke 2020) and ‘ggpubr’ (Kassambara 2020) packages for results visualization. Methods overlap (Figure 3) was plotted using the ‘UpSetR’ package (Gehlenborg 2019). To assess whether the number of sites detected as coevolving by two or more methods was significant, we defined the statistic S as the number of sites detected by at least two methods. We randomized the positions detected as coevolving independently for each method and computed S for the pseudo dataset. We repeated the procedure 10,000 times and computed a P-value as (x + 1) / 10001, where x is the number of cases where S in the randomized data sets is at least equal to the value of S on the non-randomized data.

We fitted generalized linear models for all analyzed sites independently for each type of biochemical weight used in the coevolution prediction. Whether a site was predicted as part of a coevolving group was used as a binary response variable, and the evolutionary rate of the site, its relative solvent exposure (RSA) in the protein structure, the secondary structure motif and the interaction between RSA and secondary structure were used as putative explanatory variables. As the evolutionary rate measure depends on the phylogenetic tree, we standardized them by dividing the measure by the respective family average. To account for correlated errors of sites within protein families, we further added the protein family as a random factor, resulting in a generalized linear model with mixed effects (GLMM). GLMM analyses were conducted in the R statistical software (R Core Team 2020).

The lme4 package for R (Bates et al. 2015) failed to estimate parameters for our models. We further tried different estimation procedures: (1) penalized quasi-likelihood, using the ‘glmmPQL’ function from the ‘MASS’ package (Venables and Ripley 2002), and (2) Laplace approximation, (3) adaptive Gaussian quadrature, (4) sequential reduction with the ‘glmm’ function from the ‘glmmsr’ package (Ogden 2019). All methods converged to the same model parameters, but P-values could only be obtained with the ‘glmmPQL’ method after increasing the default number of iterations. In one case (index 8), convergence was not reached after 100 iterations. We report the results from the Laplace method but used the ‘glmmPQL’ fitted models to compute variance inflation factors (VIF) using the ‘vif’ function from the ‘car’ package (Fox and Weisberg 2019), which is not compatible with the ‘glmmsr’ package.

## Data availability

All sequence alignments, phylogenies and coevolution detection results for all families are available at https://doi.org/10.6084/m9.figshare.16586915. The scripts necessary for producing the statistical analyses and figures of this manuscript are available at https://gitlab.gwdg.de/molsysevol/coevolution/atlas-of-coevolution.

## Tables

**Supplementary table S1:** Characteristics of the protein families used in the analysis.

**Supplementary table S2:** Variance inflation factors.

| variable | volume | polarity | charge | I1 | I2 | I3 | I4 | I5 | I6 | I7 | I8 |
| --- | --- | --- | --- | --- | --- | --- | --- | --- | --- | --- | --- |
| Np | 1.13 | 1.14 | 1.26 | 1.08 | 1.09 | 1.11 | 1.12 | 1.06 | 1.05 | 1.05 | 1.1 |
| Rsa | 2.58 | 2.61 | 2.82 | 2.54 | 2.54 | 2.41 | 2.54 | 2.6 | 2.7 | 2.6 | 2.57 |
| SecondaryStructure | 1.38 | 1.39 | 1.65 | 1.41 | 1.41 | 1.39 | 1.38 | 1.44 | 1.42 | 1.44 | 1.44 |
| Rsa:SecondaryStructure | 1.49 | 1.5 | 1.78 | 1.52 | 1.52 | 1.49 | 1.49 | 1.55 | 1.55 | 1.56 | 1.55 |

## Supplementary figures

**Supplementary Figure S1:** Family properties. A) Number of detected sites as a function of the number of analyzed sites. The blue line represents the best fit of a broken line regression, with the estimated breakpoint shown as a vertical orange line. B) Tree diameter as a function of median sequence length. C) Proportion of coevolving sites as a function of tree diameter.

**Supplementary Figure S3:** Effect of discretization on the estimated rate of sites. Legend as in Figure 4.

**Supplementary Figure S3:** Correlation between measures of solvent exposure. A) Principal component analysis. B) Correlogram. See Material and Methods for a description of all variables. Np.xxx: scaled evolutionary rate for property ‘xxx’. Rsa: relative solvent accessibility. NbContactX: number of residues within a distance of X Å. ResidueDepth and CalphaDepth: measures of the distance of the residue to the protein surface.

**Supplementary Figure S4:** Effect of the similarity threshold in conditional sampling. A) Conditioned on the evolutionary rate. B) Conditioned on the relative solvent accessibility (RSA).

