## Supplementary figures and images for "The structural determinants of intra-protein compensatory substitutions"

### Supplementary Figure S1

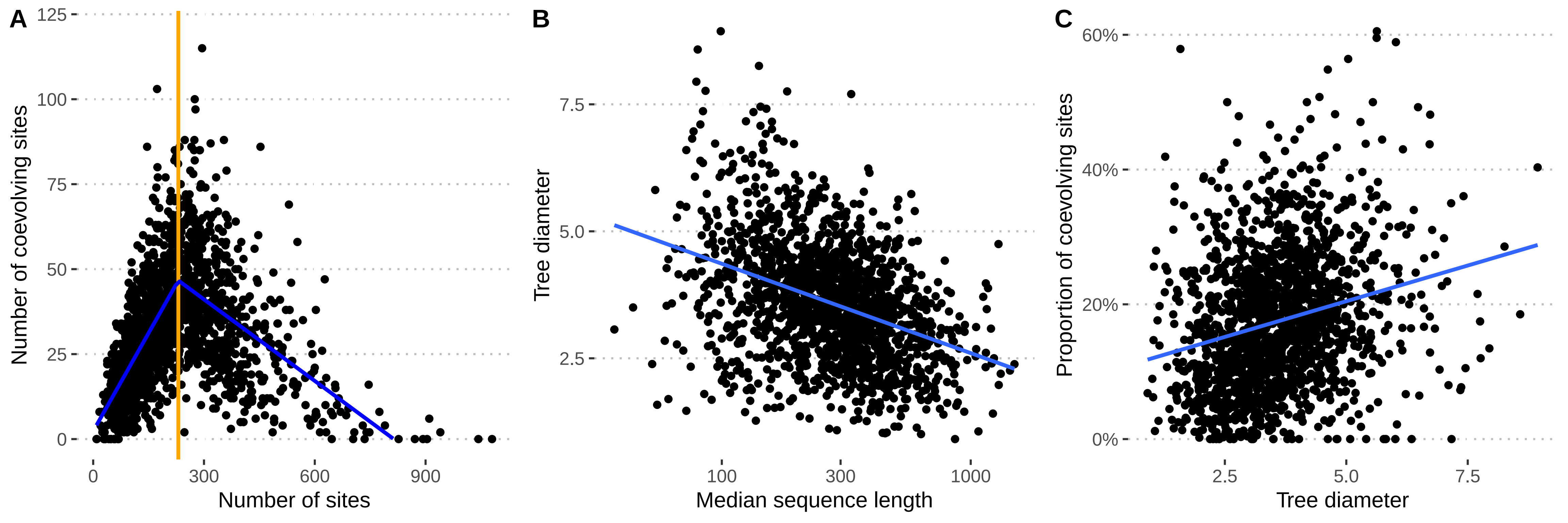

### Supplementary Figure S2

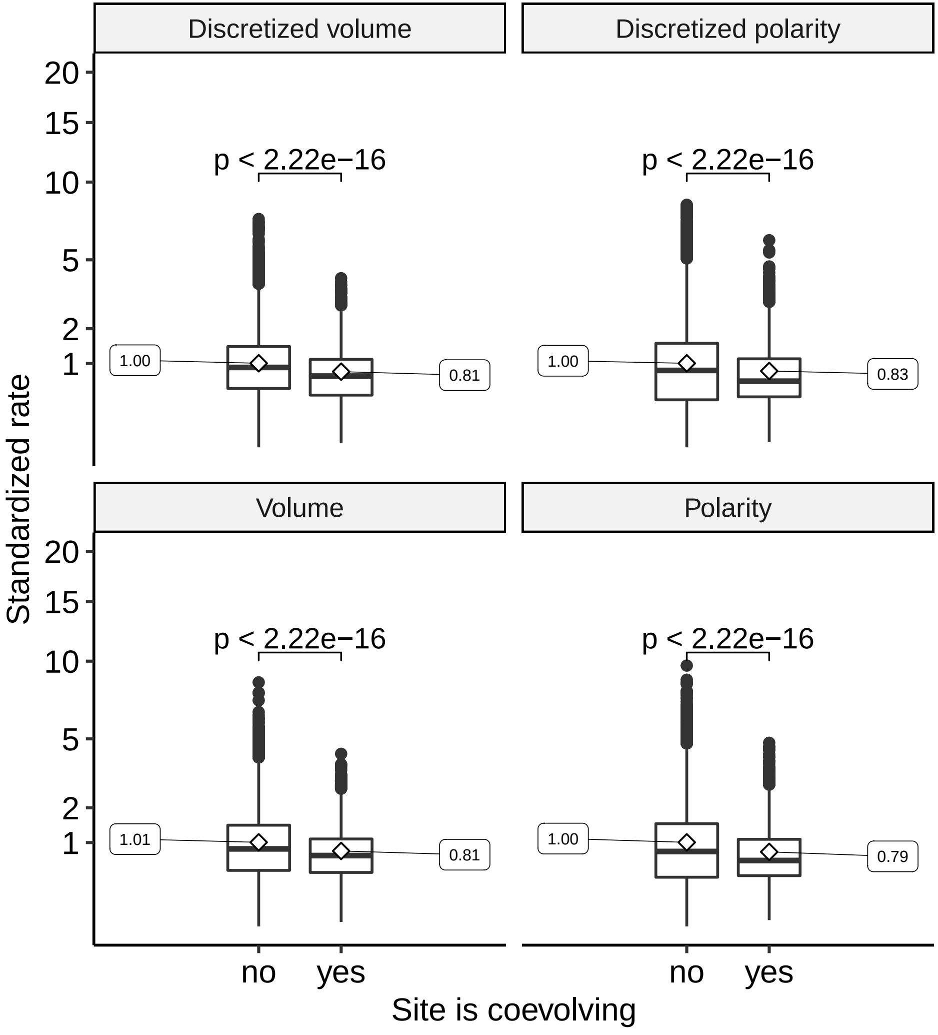

### Supplementary Figure S3

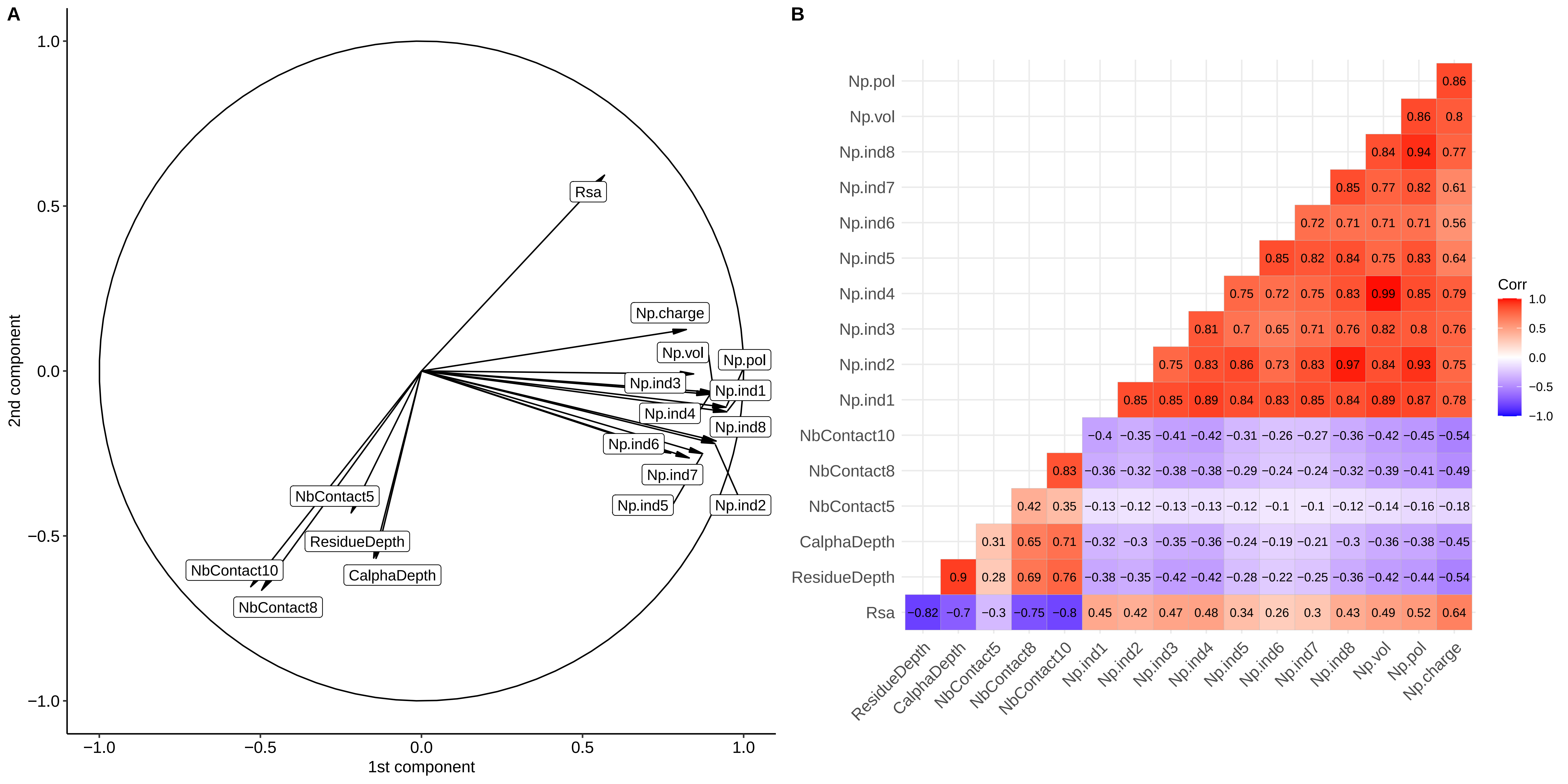

### Supplementary Figure S4

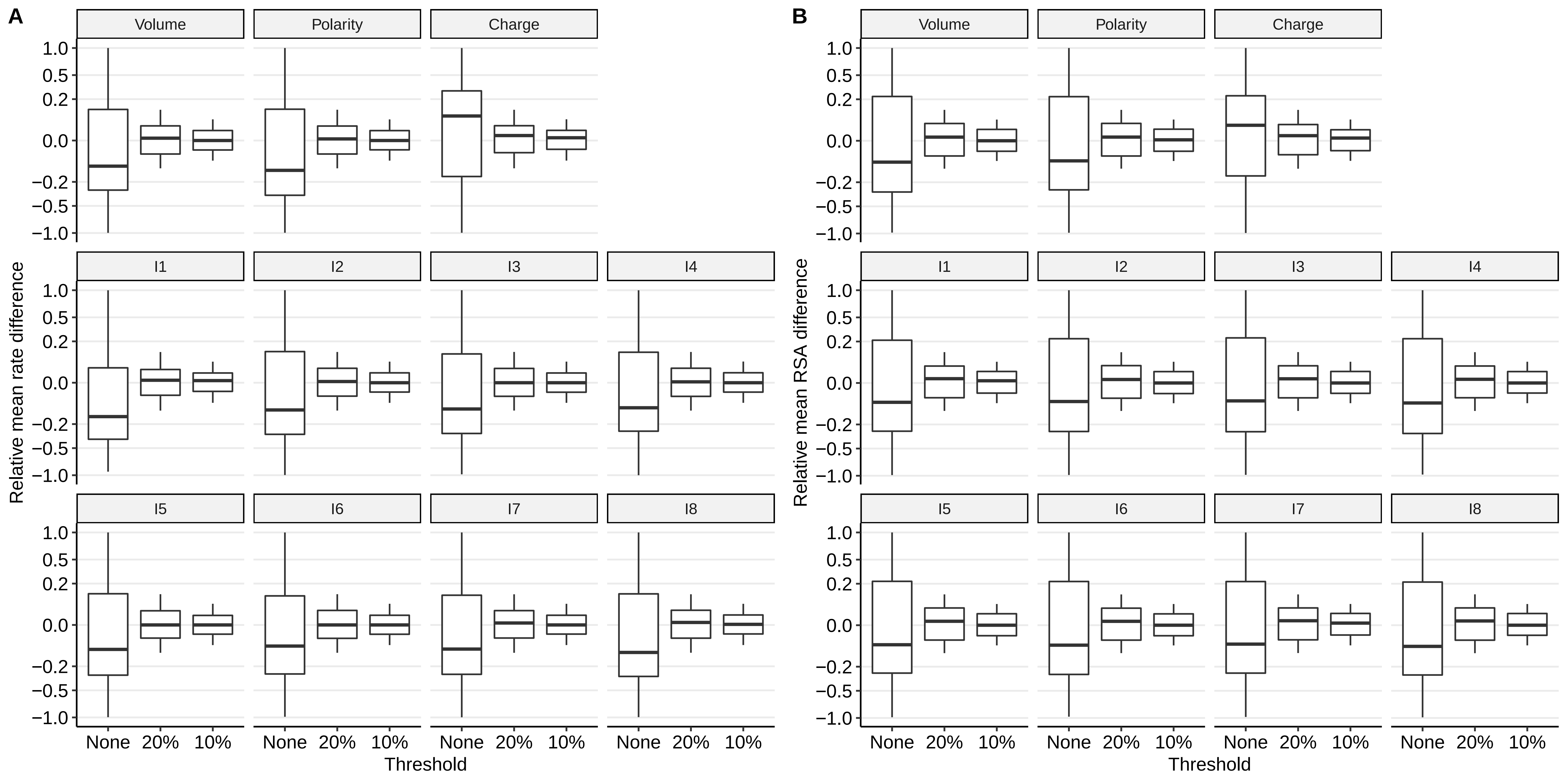
